# Evaluating Brain Flow Index from a temple-worn wearable against depth-resolved time-domain NIRS during head-down tilt and postural transitions in healthy young men

**DOI:** 10.64898/2026.09.17.751664

**Authors:** Divya Gulati, Anastasiia Rudaeva, Anirban Dutta, De’Ja Rogers, Rajveer Prajapat, Nitish Kumar, Sanchit Gupta, Deepinder Goyal, David A. Boas

## Abstract

**Significance:** Wearable optical sensors could make continuous cerebral hemodynamic monitoring practical in daily life. Because any surface head sensor also samples extracerebral tissue, demonstrating cerebral relevance requires a reference that separates deep from superficial hemodynamics.

**Aim:** We evaluated whether the Temple Brain Flow Index (Temple-BF), derived from a temple-worn photoplethysmographic wearable, covaried more strongly with the deeper (brain-assigned) than the superficial (scalp-assigned) hemodynamics recovered from time-domain near-infrared spectroscopy (TD-NIRS) during controlled postural challenges.

**Approach:** Sixty sessions from 44 healthy young adult men were analyzed across three physiological challenges: 30° head-down tilt, stand-to-squat, and stand-to-supine transitions. Optical density and mean time of flight from a three-module TD-NIRS system were inverted with a Monte Carlo-derived two-layer model to obtain scalp- and brain-layer hemoglobin time series. Temple-BF was compared with these model-derived signals using lag-adjusted and zero-lag correlations, paired brain-versus-scalp comparisons, heart rate and short-separation-HbO adjusted partial correlations.

**Results:** Median lag-adjusted correlations between Temple-BF and brain-layer ΔHbO traces were 0.912, 0.843, and 0.839 for head-down tilt, stand-to-squat, and stand-to-supine transitions, respectively; corresponding scalp-layer medians were 0.494, 0.700, and 0.767. Paired brain-minus-scalp differences were significant for head-down tilt (median 0.333, p < 0.001) and stand-to-squat (0.110, p < 0.001) but not for stand-to-supine (0.036, p = 0.082) transitions. Zero-lag brain-layer medians were 0.826, 0.701, and 0.796, and the association persisted after adjustment for heart rate (partial r = 0.832, 0.850, 0.654) and for short-separation superficial HbO (0.649, 0.570, 0.438). Transition-window correlations had medians of 0.878, 0.861, and 0.876, with directionally concordant transition responses in 59 of 60 sessions.

**Conclusions:** Temple-BF tracked model-derived brain-layer TD-NIRS ΔHbO across all three postural challenges, with larger median correlations against the brain-layer than the scalp compartment in all three protocols, and transition responses in the same direction for almost all sessions. These results support Temple-BF as a relative marker of cerebral hemodynamic change during postural perturbations. Establishing cerebral specificity at the temple, and excluding residual systemic and superficial contributions, will require flow-sensitive references and fuller systemic monitoring.

## 1 Introduction

Cerebral blood flow responds within seconds to changes in posture and exertion, and is shaped over longer timescales by sleep, diet, stress, smoking, and alcohol use^1–11^. It is increasingly regarded as a marker of vascular brain health and aging^12^. No established modality tracks cerebral hemodynamics continuously outside laboratory settings^13^. Head-worn optical wearables could fill this gap, but a sensor at the surface of the head samples scalp and systemic circulation first, so cerebral relevance must be demonstrated against a reference that separates deep from superficial tissue.

The Temple device is a small photoplethysmography (PPG) type sensor that affixes to a person’s head just over the anterior temple region. It uses a proprietary composite signal, called the Temple Brain Flow Index (Temple-BF), that aims to produce a signal indicative of cerebral blood flow. The anatomical rationale for a PPG signal over the anterior temple to be indicative of cerebral blood flow is that at the anterior temple, the frontal branch of the superficial temporal artery (a terminal external-carotid branch) meets supraorbital and supratrochlear branches of the ophthalmic artery (Fig. 1), which arises from the internal carotid^14–17^. Anatomical continuity alone, however, does not make a surface PPG signal a surrogate for cerebral blood flow. The present study therefore tests a specific prediction: if Temple-BF carries cerebrally relevant information, it should covary more strongly with the deeper, brain-assigned compartment of a depth-resolved optical reference than with the concurrently estimated superficial scalp-assigned compartment during controlled hemodynamic challenges.

**Fig. 1.**
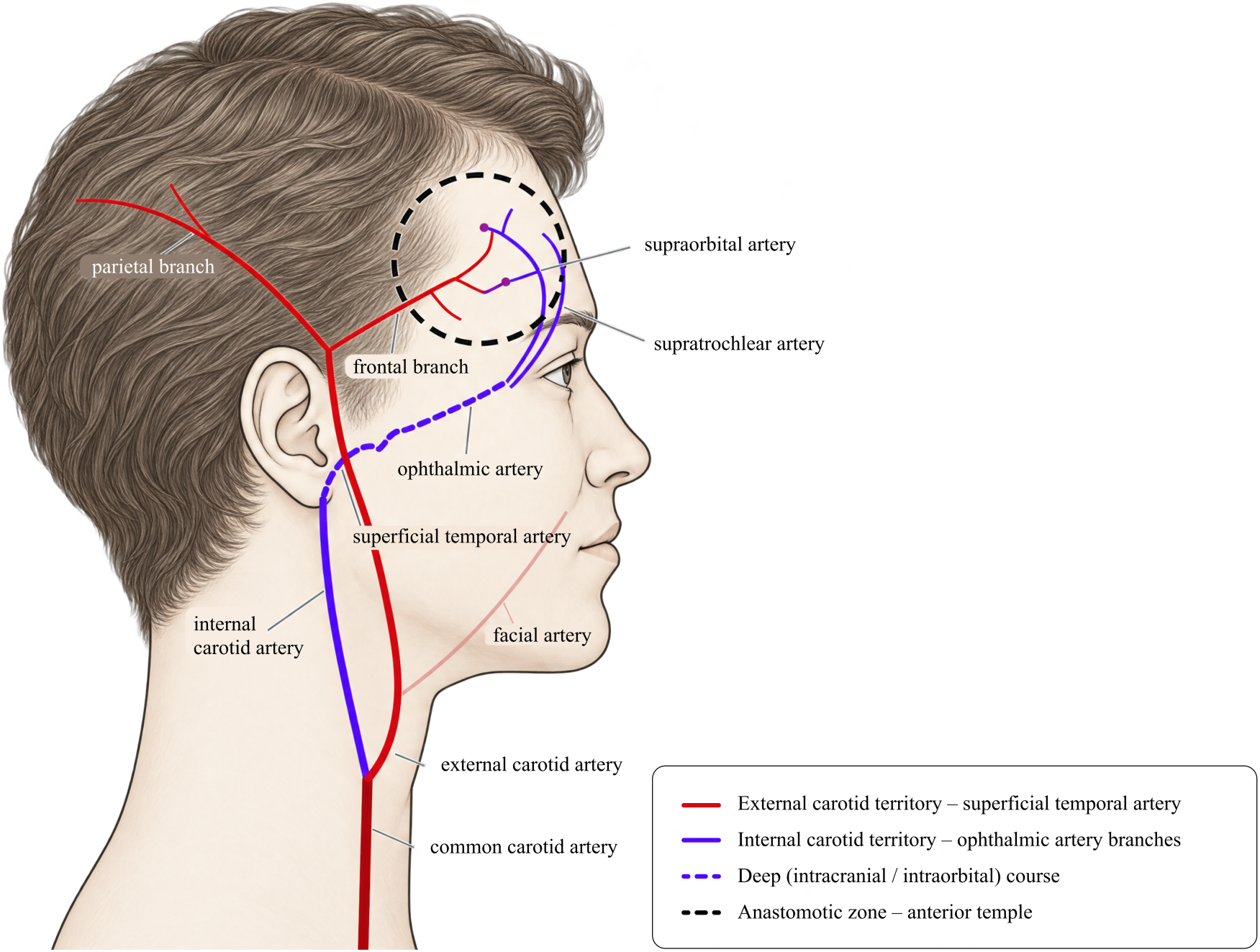
Convergence of the external and internal carotid territories at the anterior temple (schematic right lateral view, not to scale). The external carotid artery continues as the superficial temporal artery, whose frontal branch crosses the anterior temple toward the forehead. The internal carotid artery (solid) continues intracranially (dashed) and gives rise to the ophthalmic artery, whose supraorbital and supratrochlear branches resurface above the orbit; branch positions are schematic. Filled circles mark anastomoses between the frontal branch and the ophthalmic artery branches; the dashed circle marks the anastomotic zone over which the Temple sensor is worn. The face in the schematic is AI-generated.

NIRS estimates oxygenated, deoxygenated, and total hemoglobin (HbO, HbR, HbT = HbO + HbR) from multi-wavelength absorption measurements^18–20^. Changes in cerebral blood flow are accompanied by changes in hemoglobin concentrations, allowing NIRS to be an indirect measure of cerebral hemodynamic changes^21^; these are concentration estimates, not direct measurements of blood flow. Continuous-wave systems separate scalp from brain only indirectly, by combining short- and long-separation channels^22,23^. Time-domain systems additionally record the photon transit-time distribution, whose temporal moments carry depth information: longer transit times preferentially sample the brain, shorter ones the scalp^24,25^. Therefore, time-domain NIRS was selected as the comparator because it preserves photon time-of-flight information while remaining non-invasive and compatible with dynamic paradigms^18,26^. We used moments measured at three source-detector separations to recover superficial- and deeper-layer hemodynamics.

Model-derived deeper-layer ΔHbO served as the primary comparator, with ΔHbR and ΔHbT as secondary comparisons. Here, TD-NIRS was used to distinguish superficial from deeper hemodynamic responses, not as a gold-standard measure of blood flow. We induced these responses using head-tilt, squat-stand, and postural maneuvers, which are commonly used in NIRS studies to investigate hemodynamic changes^27–33^. Accordingly, we aimed to quantify the association between Temple-BF and these NIRS-derived responses and test whether it was stronger for the deeper layer than for the superficial layer.

## 2 Methods

### 2.1 Participants

This study analyzed 60 experimental sessions from 44 unique healthy young adult male participants, with 20 sessions in each protocol. Some individuals completed more than one protocol, with sessions separated by at least one week. Participants were aged 20 to 32 years (mean 24.7 ± 2.9 years) and spanned Monk Skin Tone categories 3 to 8. Exclusion criteria were a history of cardiovascular, cerebrovascular, or neurological disease; medications affecting cerebrovascular tone; hypertension; or vertigo.

The Kernel DevKit-3 (see Sec. 2.2.1) has three optical modules that are typically distributed across the forehead. To reduce potential midline vascular contamination and obtain reliable optode-skin coupling with the lateralized configuration used here, the headset was shifted over one hemisphere as shown in Fig. 2(A). The lateral placement extended toward hair-bearing regions, where hair can attenuate NIRS signals and degrade coupling^20^. Recruitment was restricted to men in this study as a pragmatic attempt to reduce hair-related measurement artefacts.

**Fig. 2.**
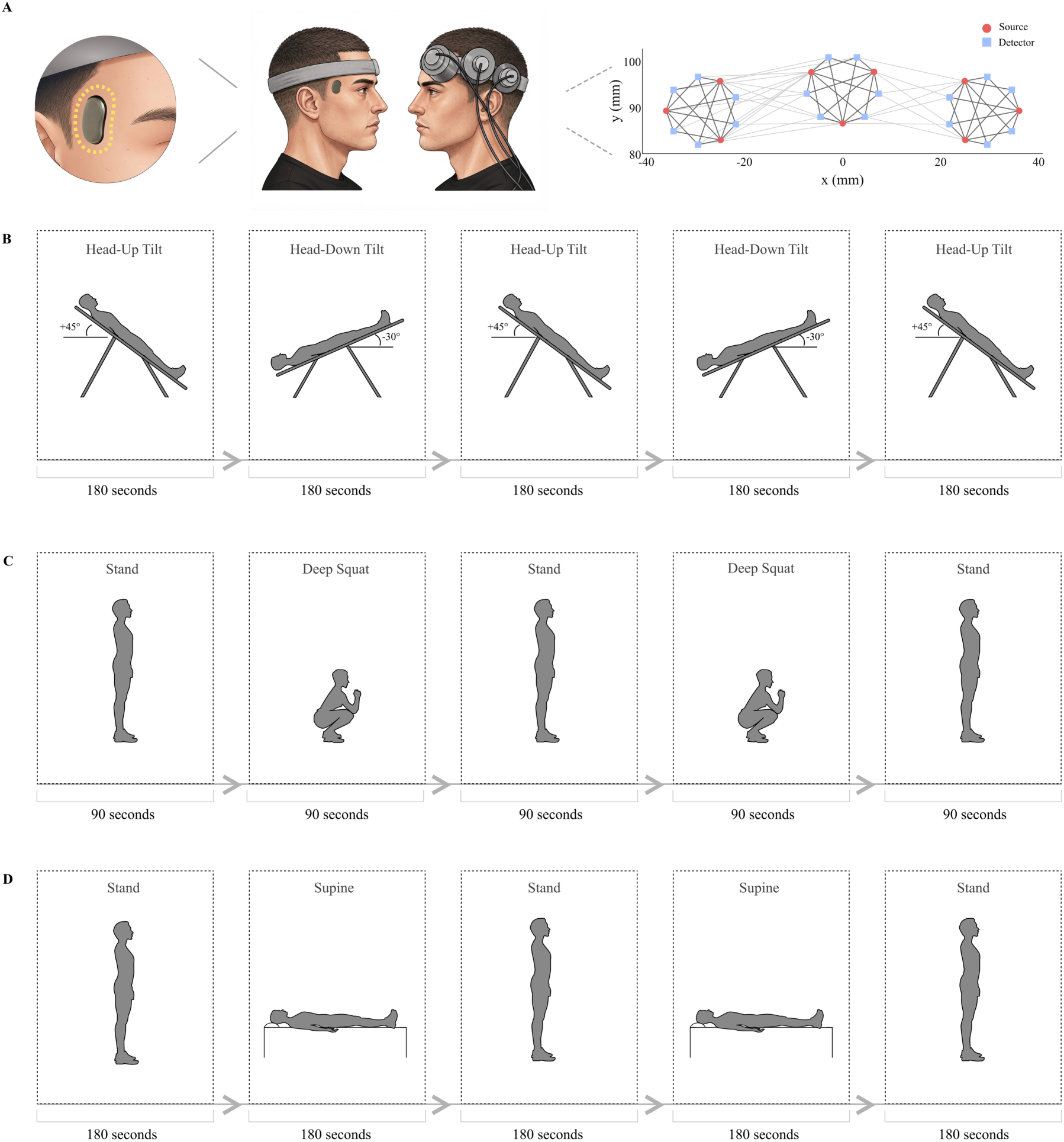
Schematic representation of the experimental setup and block design for the different protocols. (A, left) A participant wearing the Temple and Kernel NIRS devices during a recording session. This schematic was enhanced using AI. (A, right) The module layout, with each module having 3 sources and 6 detectors, and the number of remaining channels after post-pruning for a representative subject; each source-detector pair forms a channel (gray), and dark gray lines indicate channels with a source-detector separation greater than 25 mm. (B) Head-down tilt, (C) stand-to-squat, and (D) stand-to-supine protocols, each consisting of five sequential blocks of the indicated duration. Each block is represented by the dashed box, and the gap between blocks indicates the transition period between the blocks.

### 2.2 Experimental setup

#### 2.2.1 Devices

We used the Kernel DevKit-3 TD-NIRS system, which consists of three modules, each containing 6 detectors and 3 sources, two wavelengths (690 and 905nm) and samples at 15.89Hz.

The Temple device is a temple-worn wearable combining multi-wavelength reflectance photoplethysmography (PPG), heart rate, accelerometry, and temperature sensing. The PPG sensor uses green, red, and infrared illumination. Different wavelengths have overlapping but wavelength-dependent sampling of skin and subdermal vasculature, with longer wavelengths often sampling deeper on average^34–36^. Temple-BF is a proprietary composite index integrating (i) an AC/DC perfusion-related PPG feature, (ii) relative hemoglobin-related optical features, (iii) pulse-waveform morphology, heart rate features, and body temperature dynamics, and (iv) accelerometer information used for artefact handling and contextualization. Temple-BF was exported at 1 Hz using algorithm version 10.0. The Temple-BF algorithm was fixed prior to the commencement of this study; no tuning or optimization was performed on the algorithm based on the data collected for this study.

#### 2.2.2 Recording session setup

Recordings took place in a room at approximately 25°C, with the lights switched off and recording screens facing away from participants. Sessions were scheduled between 09:00 and 19:30h. Alcohol and caffeine were prohibited for the 12h preceding each session, and the final meal was completed no less than 1h before recording. Written informed consent was obtained before study procedures. Participants then completed a 5–7min seated rest period while task instructions were reviewed. NIRS and Temple data were acquired simultaneously. The NIRS module was positioned over the forehead and the Temple device over the contralateral anterior temple, with lateral separation intended to minimize optical crosstalk [Fig. 2(A)]. The NIRS assembly was secured using the commercial-product headband configuration.

### 2.3 Study design

Each recording session consisted of one of three experimental protocols, a head-down tilt protocol, a stand-to-squat protocol, or a stand-to-supine orthostatic transition protocol, as shown in Fig. 2. For all protocols, participants held each position quietly for the duration of the block, refraining from speech, jaw or eyebrow movement, and postural adjustment. In the head-down tilt protocol Fig. 2(B), participants were positioned on an inversion table, which was moved by the investigator to achieve the required body positions. The protocol consisted of five consecutive blocks: 45° head-up tilt, 30° head-down tilt (−30° in the convention used in the tilt literature), 45° head-up tilt, 30° head-down tilt, and 45° head-up tilt. Each block lasted 180s. In the stand-to-squat protocol, Fig. 2(C), participants completed five consecutive blocks in the following order: standing, deep squat, standing, deep squat, and standing. Each block lasted for 90s. During the squat blocks, participants were instructed to squat as low as possible while keeping their feet no wider than shoulder-width apart. If participants were unable to maintain the squat position comfortably, they were allowed to gently touch their hips to the support behind them but were instructed not to transfer their body weight onto it. In the stand-to-supine orthostatic transition protocol, Fig. 2(D), participants completed five consecutive blocks: standing, supine lying, standing, supine lying, and standing. The supine position was performed on a bed, and each block lasted 180s. For both the stand-to-squat and stand-to-supine orthostatic transition protocols, participants were instructed to transition gently between positions. Event timings for each protocol were saved in an events file and used for alignment and epoch definition.

### 2.4 TD-NIRS processing and two-layer model estimation

NIRS data were exported from Kernel’s server as Moment SNIRF files^37^. Before export, raw time-domain photon-count data underwent Kernel-side preprocessing, including trimming of recording edges, removal of channels classified as low quality, histogram noise-floor correction, and conversion of photon arrival-time histograms into moment-based features. The exported files contained intensity, mean time of flight (TOF), and variance of TOF for each source-detector channel and wavelength.

Signal quality was evaluated on the exported intensity data using Cedalion^38^ v26.5.1.dev2. Scalp coupling index and peak spectral power were computed over 5s windows for each channel. A channel entered the good-channel mask only when at least 60% of windows passed both criteria, using thresholds of 0.5 for scalp coupling index and 0.05 for peak spectral power. This mask restricted all subsequent fitting to retained channels. Source-detector distances were computed from the SNIRF optode geometry and used to define module-wise channel groups and separation constraints.

Intensity was converted to optical density (OD) using a log-ratio transformation; OD and mean TOF served as the observables for the inversion. Temperature regression was applied to mean TOF to correct for the temperature dependence of the instrument response function. For each channel, the corresponding source-temperature trace was used to fit TOF(t) = aT(t) + b, and the fitted component was subtracted to obtain TOF_corr(t) = TOF(t) − [aT(t) + b]. Temperature regression was applied to mean TOF only, because photon timing is referenced to the instrument response and is susceptible to thermal timing drift, whereas OD is a within-channel log-ratio of intensity and is not similarly affected. Two preprocessing variants were then evaluated. The primary analysis used OD and temperature-corrected TOF without additional motion correction. In a secondary sensitivity analysis, shown in the supplementary figures, temporal derivative distribution repair (TDDR) motion correction^39^ was applied to both OD and temperature-corrected TOF before model fitting. For both variants, OD and TOF were low-pass filtered at 0.5Hz and baseline-zeroed by subtracting the median of the first 60s before inversion.

To estimate absorption changes in superficial and deeper layers from the OD and TOF time traces, layer-specific sensitivity terms were required. These sensitivities depend on measurement geometry, modeled head anatomy, and baseline optical properties. The source-detector geometry was defined by the arrangement of 3 sources and 6 detectors per Kernel module. Measurements were modeled using a semi-infinite two-layer medium, with an 8 mm superficial layer representing scalp and skull, and the remaining semi-infinite layer representing the deeper compartment. Baseline optical properties were estimated separately for each subject, module, and wavelength, 690 and 905nm, from the absolute moment data using a nonlinear fit to the homogeneous diffusion equation. Temperature correction was disabled during this stage; instead, unknown instrument response function (IRF) moments were included in the nonlinear fitting algorithm and estimated simultaneously with the tissue optical properties. This fit provided subject- and module-specific wavelength-dependent absorption and reduced scattering coefficients.

These optical properties were then used in Monte Carlo simulations of the two-layer geometry to estimate the sensitivity of each channel to absorption changes in the superficial and deeper layers, following the moment-sensitivity framework^24,26^. The superficial and deeper layers used the same reduced scattering coefficient estimated for the corresponding subject and module. The superficial-layer absorption coefficient was given by the homogeneous fit, while the deeper-layer absorption coefficient was set to 1.5× the superficial-layer value to account for the greater total hemoglobin concentration in the brain compared with the scalp. Monte Carlo simulations were performed using Monte Carlo eXtreme for OpenCL (MCXCL)^40,41^.

Each simulation was run in two chunks of 5 × 10⁸ photons, with up to 5 × 10⁶ detected photons retained per chunk; detected pathlength histories were combined during post-processing. Throughout this manuscript, “brain layer” denotes this model-assigned deeper compartment and not anatomically segmented or directly observed cortex.

Combined photon histories were used to compute moment-based sensitivity terms for each source-detector separation and wavelength. Mean partial pathlength (MPP) describes the sensitivity of OD to absorption change through the average distance traveled within each modeled layer, and the mean-time sensitivity factor (MTSF) describes the sensitivity of mean photon arrival time to absorption change in each layer^24,25^. A variance sensitivity factor (VSF), describing sensitivity of the arrival-time variance, was also computed but was not used in the final inversion; only MPP and MTSF were carried forward.

For each subject and module, retained within-module channels were fitted together using weighted linear least squares. MCX-derived sensitivities were interpolated onto the actual source-detector distances of the retained channels. Optical density changes were paired with MPP, and mean-TOF changes were paired with MTSF, consistent with the moment-sensitivity framework^24^. For each wavelength and time point, the model estimated superficial and deeper-layer absorption changes as

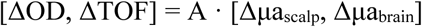

where A contains the MPP and MTSF sensitivity terms. Channel weights were set as w = 1/σ, with σ estimated from the standard deviation of temporal differences in the signal, so that noisier channels received lower weight.

A homogeneous fit was computed on the same retained channels for comparison. In this case, the superficial and deeper-layer sensitivities were summed into a single effective sensitivity per channel, and the model solved for one absorption-change parameter per wavelength and time point. This homogeneous estimate represents the absorption change that would be recovered if the measured OD and TOF changes were attributed to a single effective tissue compartment, and is distinct from the homogeneous diffusion fit used earlier to estimate baseline optical properties.

The inversion produced wavelength-specific absorption changes at 690 and 905nm, which were converted to hemoglobin concentration changes using the Prahl extinction coefficients^42^ by solving Δμa = E · Δc with the pseudoinverse of the extinction matrix, Δc = E⁺ · Δμa. This produced ΔHbO and ΔHbR traces in micromolar units for the homogeneous, superficial, and deeper-layer estimates, with ΔHbT computed as ΔHbO + ΔHbR.

Modules were screened using conditioning metrics derived from the two-layer inversion. For each subject, module, protocol, and wavelength, the weighted sensitivity matrix was reconstructed using the same rows used in the inversion: OD rows contained MPP sensitivities, TOF rows contained MTSF sensitivities, and the two columns represented superficial-layer and deeper-layer absorption changes.

Three metrics were calculated from this weighted sensitivity matrix. First, the condition number was calculated as the ratio of the largest to smallest singular value of the matrix. Larger condition numbers indicate that superficial and deeper-layer absorption changes are less stably separable from the OD and TOF measurements. Second, the deeper-layer inverse gain was calculated from the deeper-layer row of the pseudoinverse matrix, with larger values indicating greater amplification of measurement noise into the estimated deeper-layer absorption change. Third, the weighted deeper-layer column fraction was calculated as the relative norm of the deeper-layer sensitivity column compared with the total superficial-plus-deeper sensitivity column norm, with smaller values indicating that the measurement was dominated by superficial sensitivity.

For each subject-module-protocol combination, the larger condition number, larger deeper-layer inverse gain, and smaller weighted deeper-layer column fraction across 690 and 905nm were used for screening. A module was retained if the larger of its two wavelength-specific condition numbers and the larger of its two wavelength-specific deeper-layer inverse gains were each below the 80th percentile of the pooled distribution across all subject-module-protocol combinations, and if the smaller of its two wavelength-specific weighted deeper-layer column fractions was at least 0.08. These thresholds were chosen as dataset-relative inversion-stability criteria to exclude modules with the poorest conditioning or very weak deeper-layer sensitivity. Passing modules were averaged within each session to obtain homogeneous, superficial, and deeper-layer ΔHbO, ΔHbR, and ΔHbT traces.

For comparison with conventional OD-derived hemoglobin signals, retained channels were also converted from intensity to OD and then to hemoglobin concentration changes using the modified Beer-Lambert law. In this analysis, OD was bandpass filtered between 0.001 and 2.0Hz before conversion. The conversion used a differential pathlength factor of 6.0 and the Prahl wavelength-dependent extinction coefficients. Short-SDS and long-SDS OD-derived traces were obtained by averaging retained channels with source-detector separations ≤9 mm and >9 mm, respectively.

### 2.5 Temple data processing

Temple-BF and heart rate values were exported from the Temple application at 1 Hz. Event timings were parsed from the protocol event files, and Temple data were aligned to NIRS using SNIRF recording-start metadata timestamps. For visualization and correlation analyses, Temple-BF and heart rate traces were interpolated onto the same 1Hz time grid used for the NIRS traces.

### 2.6 Visualization and statistical analysis

Only modules passing the inversion-conditioning screen entered downstream analysis. Passing modules were averaged within each session to obtain session-level homogeneous, superficial, and brain-layer traces; group traces were then obtained by averaging session-level traces within protocol. Unless otherwise specified, time series were smoothed with a forward-backward exponential moving-average filter (α = 0.35), applied symmetrically to avoid phase delay. Because the traces were sampled at 1s intervals, α = 0.35 corresponds to an approximate single-pass EMA time constant of 2.3s. All inferential analyses were protocol-specific, and no pooled model combined sessions across protocols; therefore, cross-protocol within-person dependence was not treated as independent evidence in a single pooled test.

Temporal correspondence was quantified with lagged Pearson correlations. The primary analysis used model-derived brain-layer ΔHbO, with superficial-layer ΔHbO analyzed in parallel. NIRS and Temple-BF were interpolated onto a common 1 Hz grid over the shared event-bounded interval. For each session, Pearson r was evaluated at every integer lag within a ±30 s window, and the largest coefficient defined the lag-adjusted correlation. Under our sign convention, a positive optimal lag means the NIRS signal led Temple-BF. Group-level analysis used one-sided Wilcoxon signed-rank tests against zero^43^. Session-level brain-layer and scalp-layer lag-adjusted correlations were compared with one-sided paired Wilcoxon signed-rank tests of the hypothesis that the brain-layer correlation was larger.

Zero-lag Pearson correlations were also calculated between brain-layer ΔHbO and Temple-BF without temporal shifting. To assess whether heart rate variation accounted for the observed association, first-order partial correlations controlled for Temple-derived heart rate. Brain-layer ΔHbO and Temple-BF were aligned using the session-specific optimal lag from the lagged analysis, and heart rate was shifted by the same lag. This adjustment controls only heart rate and does not remove other systemic drivers such as arterial blood pressure, arterial CO₂, respiration, cardiac output, or superficial vasomotion^44^.

As an additional sensitivity analysis for superficial contributions, we repeated the partial-correlation analysis while controlling for short-separation NIRS-HbO. For each session, HbO traces were recomputed from retained short source-detector channels using intensity-to-OD conversion described in Sec. 2.4. Short-separation HbO traces were then averaged across retained channels with source-detector separation ≤9 mm and used as a superficial nuisance regressor. Model-derived brain-layer ΔHbO and Temple-BF traces were aligned using the same session-specific optimal lag from the primary lagged-correlation analysis. We then calculated the partial correlation between brain-layer ΔHbO and Temple-BF after regressing the short-separation HbO trace from both signals. This analysis tested whether the association between Temple-BF and model-derived brain-layer ΔHbO persisted after removing variance shared with superficial short-separation HbO.

Transition analyses quantified baseline-to-task responses. Each full protocol trace was z-scored separately for each signal, and predefined windows were extracted around baseline-to-task transitions: 90s for head-down tilt, 45s for squat, and 90s for stand-to-supine. For each transition, response magnitude was calculated as the post-transition mean minus the pre-transition mean. Repeated transitions of the same type were averaged within the session to yield one NIRS value and one Temple-BF value per protocol session. One-sided Wilcoxon signed-rank tests against zero were used, and whether the two transition changes shared the same sign was summarized within each session.

Temporal correlations were also calculated within baseline-to-task transition windows between z-scored brain-layer ΔHbO and Temple-BF. Repeated transition coefficients were Fisher r-to-z transformed, averaged within session, and transformed back to r. Windows were labeled as meeting a predefined criterion when r > 0.5 and the Pearson p value was <0.01. As a supplementary analysis, transition-window correlations were also shown without Fisher transformation or within-session averaging. In this, each repeated baseline-to-task transition window was retained as a separate observation, providing the distribution of individual transition-window correlations rather than one averaged value per session.

Conventional Bland-Altman analysis is intended to assess agreement between methods measuring the same quantity and requires clinically or scientifically meaningful limits of acceptable disagreement^45–47^. Because NIRS-ΔHbO (micromolar) and Temple-BF (dimensionless) share no physical unit, each trace was first z-scored within the session; paired transition differences (Temple-BF minus NIRS-ΔHbO) and mean ± 1.96 SD limits were then calculated. Z-scoring imposes comparable location and scale and can move the mean standardized difference toward zero; therefore, these plots were interpreted as descriptive agreement in standardized transition-response magnitude rather than agreement in absolute measurement units.

## 3 Results

Simultaneous NIRS and Temple recordings were collected while participants performed one of three physiological challenge protocols (Fig. 2). Each session consisted of five sequential blocks involving repeated posture or tilt transitions. The head-down tilt protocol alternated between 45° head-up tilt and 30° head-down tilt. The stand-to-squat protocol alternated between standing and deep squatting. The stand-to-supine orthostatic transition protocol alternated between standing and supine rest.

### 3.1 Depth-resolved hemoglobin responses

Hemoglobin responses were estimated with the two-layer time-domain model, which returned superficial and deeper “brain-layer” absorption changes at each time point, together with a homogeneous single-compartment fit on the same channels. Modules were screened for inversion conditioning before averaging. All 60 sessions retained at least one module after screening, with a median of two passing modules per session across protocols.

Fig. 3 shows group-averaged homogeneous, superficial, and brain-layer estimates. Transitions into head-down tilt, deep squat, or supine posture produced block-locked changes in each model output. ΔHbO and ΔHbT generally increased during task blocks and moved toward baseline after the reverse transition, whereas ΔHbR showed an opposite or weaker pattern. Model-derived amplitudes differed across compartments. The brain-layer ΔHbO response increased gradually within the task blocks rather than changing instantaneously, reaching approximately half of its within-block peak within the first 10 to 20s and reaching its peak response after roughly 60 to 90s. Because the NIRS modules were fixed tightly to the head, we did not expect the probes to move substantially relative to the scalp during the posture and tilt transitions. Therefore, motion correction was not included in the primary preprocessing pipeline. However, to test whether the results were sensitive to residual motion-related artefacts, we also repeated the same depth-resolved model pipeline after applying TDDR motion correction to both OD and TOF before inversion (Fig. S1). The overall block-locked response structure was preserved, but response amplitudes were smaller after TDDR preprocessing, with the strongest attenuation in the superficial-layer traces, especially during head-down tilt. The HDT brain-layer ΔHbR trace also differed between preprocessing variants, showing an opposite-going pattern after TDDR compared with the analysis without TDDR. All subsequent analyses are reported without TDDR.

**Fig. 3.**
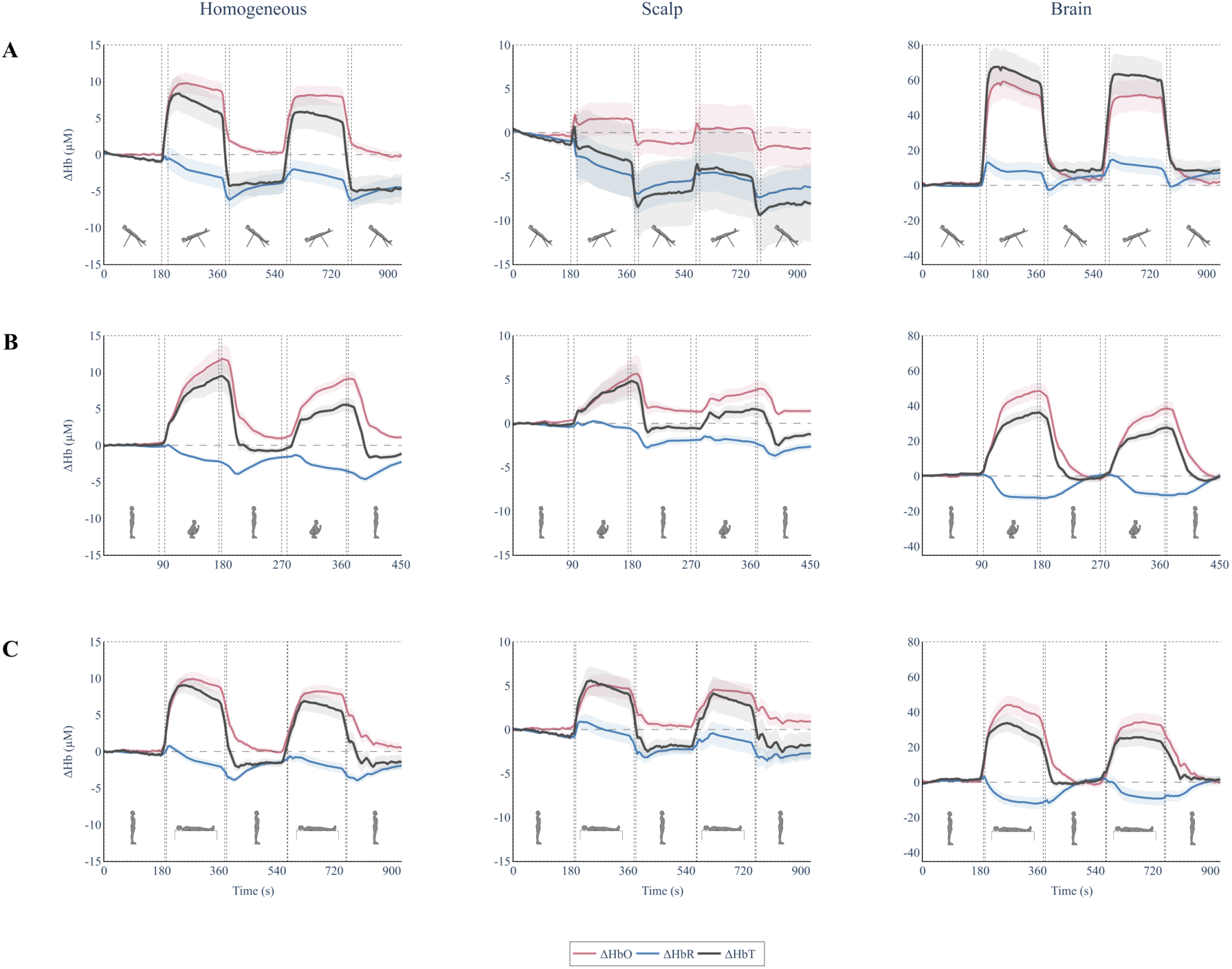
Model-derived NIRS hemoglobin responses across the three physiological challenge protocols (n = 20 sessions per protocol). Group-averaged hemoglobin traces are shown for (A) head-down tilt, (B) stand-to-squat, and (C) the stand-to-supine protocol. Each row corresponds to one protocol, and the three columns show the homogeneous estimate, the scalp estimate, and the brain estimate from the two-layer model. Each protocol consisted of five blocks, indicated by the dashed boxes, with the gaps between them indicating the transition periods between consecutive blocks. Traces show model-derived ΔHbO (pink), ΔHbR (blue), and ΔHbT (dark gray), with shaded bands representing SEM across sessions. Before averaging, modules were screened using inversion-conditioning criteria, and only passing modules were averaged within each session. Note the differing y-scale across columns.

### 3.2 Temple-BF shows larger median correlations with brain-layer than superficial-forehead ΔHbO

Across all three physiological challenge protocols, Temple-BF closely matched the temporal pattern observed for model-derived NIRS-ΔHbO. It was higher when participants were in the head-down tilt, deep squat, and supine positions compared with the head-up tilt or standing positions (Fig. 4).

**Fig. 4.**
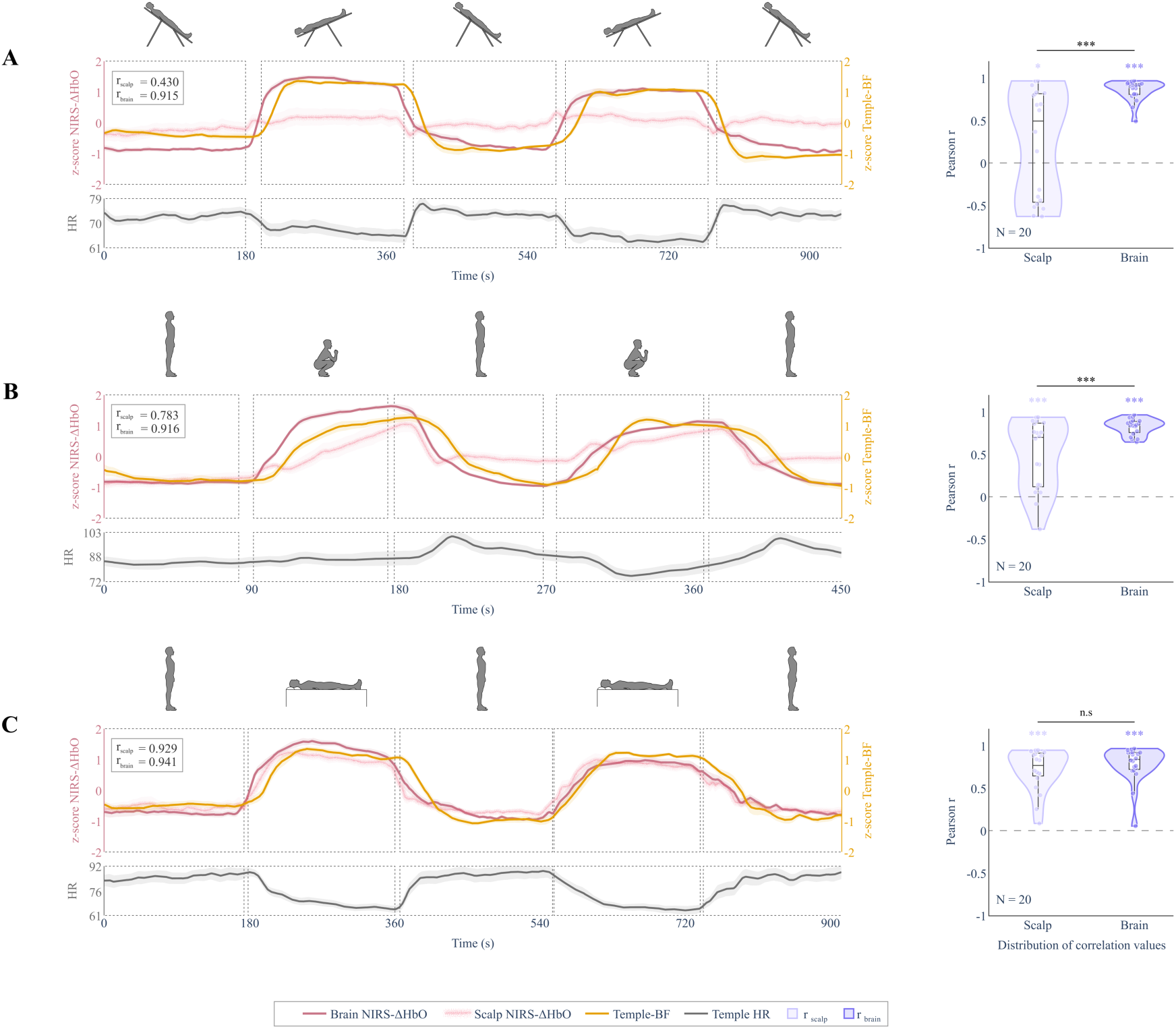
Full-session temporal correspondence between model-derived NIRS-ΔHbO and Temple-BF across sessions and protocols (n = 20 sessions per protocol). For each protocol, group-averaged z-scored time waveforms are shown across the full recording duration for (A) head-down tilt, (B) stand-to-squat, and (C) the stand-to-supine protocol. Brain ΔHbO is shown as a dark-pink trace, scalp ΔHbO as a light-pink trace, and Temple-BF in gold. Temple-derived heart rate is shown in gray below each panel. Dashed boxes indicate the task blocks, with the gaps between them indicating transition periods. The r_scalp_ and r_brain_ values annotated on the time-trace panels are correlations computed on the group-mean traces; the violin plots beside them show the session-level distributions. Violin plots show the distribution of session-level lag-adjusted Pearson correlation coefficients, separately for the scalp and brain estimates. Significance relative to zero was evaluated using a one-sided Wilcoxon signed-rank test; “***” indicates p < 0.001 and “*” indicates p < 0.05. Brackets above the scalp and brain violin plots show the paired brain-minus-scalp comparison within each protocol, tested using a one-sided paired Wilcoxon signed-rank test.

To quantify the temporal correspondence, session-level lag-adjusted Pearson correlations were calculated between Temple-BF and model-derived NIRS-ΔHbO over lags from −30 to +30s. The primary comparison used the brain-layer ΔHbO estimate, while the superficial-layer estimate was analyzed in parallel. As shown in the violin plots on the right side of Fig. 4, median lag-adjusted correlations between Temple-BF and brain-layer ΔHbO were 0.912 for head-down tilt, 0.843 for stand-to-squat, and 0.839 for stand-to-supine protocols. These correlations were significantly greater than zero in all three protocols using one-sided Wilcoxon signed-rank tests (all p < 0.001).

Correlations with the scalp-layer ΔHbO estimate were lower, with median values of 0.494 for head-down tilt, 0.700 for stand-to-squat, and 0.767 for stand-to-supine protocols. Although the scalp-layer correlations were also positive, paired brain-minus-scalp comparisons showed that brain-layer ΔHbO correlations were significantly higher than scalp ΔHbO correlations for head-down tilt (median paired difference = 0.333, p < 0.001) and stand-to-squat (median paired difference = 0.110, p < 0.001), but not for stand-to-supine (median paired difference = 0.036, p = 0.082). Full-session results are summarized in Table 1.

**Table 1.** Full-session correspondence between Temple-BF and model-derived NIRS-ΔHbO by protocol (n = 20 sessions per protocol). Values are session-level medians. Brain and scalp columns show maximum lag-adjusted Pearson correlations between Temple-BF and the corresponding modeled compartment. The paired brain-minus-scalp column reports the session-level paired difference between brain-layer and scalp lag-adjusted correlations, with p values from one-sided paired Wilcoxon signed-rank tests testing whether brain-layer correlations were greater than scalp correlations. Zero-lag correlations are shown for the brain layer estimate without lag adjustment. Partial r values are brain-layer partial correlations controlling separately for Temple-derived heart rate or OD-derived short-SDS HbO. All other p values are from one-sided Wilcoxon signed-rank tests against zero.

| Protocol | Lag-adjusted brain-layer ΔHbO r | Lag-adjusted scalp ΔHbO r | Paired brain – scalp ΔHbO r | Zero-lag brain-layer r | Partial brain-layer r (HR) | Partial brain-layer r (short-SDS HbO) |
| --- | --- | --- | --- | --- | --- | --- |
| Head-down tilt | 0.912, p < 0.001 | 0.494, p = 0.027 | 0.333, p < 0.001 | 0.826, p < 0.001 | 0.832, p < 0.001 | 0.649, p < 0.001 |
| Stand-to-squat | 0.843, p < 0.001 | 0.700, p < 0.001 | 0.110, p < 0.001 | 0.701, p < 0.001 | 0.850, p < 0.001 | 0.570, p < 0.001 |
| Stand-to-supine | 0.839, p < 0.001 | 0.767, p < 0.001 | 0.036, p = 0.082 | 0.796, p < 0.001 | 0.654, p < 0.001 | 0.438, p < 0.001 |

The optimal lags for the brain-layer ΔHbO correlations were consistently positive across protocols, with median lags of 13 to 15 s, whereas scalp ΔHbO lags were more variable, especially during head-down tilt. Positive lags indicate that Temple-BF followed NIRS-ΔHbO; but these lags should be interpreted as descriptive alignment parameters rather than physiological delay times.

Temple-derived heart rate is shown below each corresponding trace panel as a physiological reference. Across protocols, heart rate generally decreased during periods when Temple-BF increased, showing an opposite pattern.

### 3.3 Chromophore dependence

The same lagged-correlation analysis was applied to ΔHbR and ΔHbT as supplementary analyses. ΔHbR showed a weaker and less consistent relationship with Temple-BF than ΔHbO (Fig. S2). For ΔHbR, scalp correlations were positive across protocols, and the brain estimate was positive for head-down tilt, but brain ΔHbR was not positively associated with Temple-BF during stand-to-squat or stand-to-supine. In contrast, ΔHbT showed positive correlations with Temple-BF across all three protocols (Fig. S3). Median brain-layer ΔHbT correlations were 0.915 for head-down tilt, 0.843 for stand-to-squat, and 0.850 for stand-to-supine, with all three significantly greater than zero. Overall, the Temple-BF association was strongest and most consistent for ΔHbO and ΔHbT, whereas ΔHbR showed protocol- and compartment-dependent behavior.

### 3.4 Zero-lag, heart rate and short channel derived HbO-adjusted sensitivity analyses

Zero-lag Pearson correlations were calculated as a simpler, non-lag-adjusted measure of the temporal relationship between brain-layer ΔHbO and Temple-BF. Median smoothed zero-lag correlations remained positive across all three protocols: 0.826 for head-down tilt, 0.701 for stand-to-squat, and 0.796 for stand-to-supine, with one-sided Wilcoxon signed-rank tests showing p < 0.001 for all protocols (Table 1). Although these values were lower than the lag-adjusted correlations, they remained strongly positive, indicating that the observed temporal correspondence was not created only by selecting a favorable temporal shift.

Heart rate-adjusted partial correlations were calculated to test whether changes in heart rate alone could explain the observed relationship between Temple-BF and brain-layer NIRS-ΔHbO. Median brain-layer partial correlations after controlling for Temple-derived heart rate remained positive: 0.832 for head-down tilt, 0.850 for stand-to-squat, and 0.654 for stand-to-supine, with all three distributions significantly greater than zero (p < 0.001, Table 1). This indicates that the association between Temple-BF and brain-layer NIRS-ΔHbO was not eliminated by linear adjustment for heart rate alone.

Fig. S4 compares Temple-BF, model-derived brain and scalp ΔHbO, and conventional OD-derived ΔHbO from short- and long-SDS channels. The violin plots summarize the session-level lag-adjusted correlations between Temple-BF and OD-derived ΔHbO from short- and long-SDS channels, showing that both channel groups carried Temple-BF-related temporal structure. Short-SDS OD-derived HbO was used as a superficial-sensitivity regressor because channels with source-detector separations ≤9 mm are expected to be dominated by scalp and extracerebral hemodynamics. This analysis tested whether the Temple-BF association with brain-layer NIRS-ΔHbO remained after accounting for the component linearly shared with the short-SDS HbO signal. After controlling for OD-derived short-SDS HbO, the median partial correlations between brain-layer ΔHbO and Temple-BF were reduced relative to the original lag-adjusted correlations but remained positive across all three protocols: 0.649 for head-down tilt, 0.570 for stand-to-squat, and 0.438 for stand-to-supine (Table 1). These partial correlations were also significantly greater than zero. Together, these results suggest that part of the Temple-BF-related temporal structure overlapped with OD-derived channel-level HbO representative of the scalp signal, but the relationship between Temple-BF and the model-derived brain-layer ΔHbO was not explained entirely by the short-SDS HbO signal.

### 3.5 Baseline-to-task transition responses

To evaluate the direction and consistency of task-related changes, baseline-to-task transition responses were calculated from z-scored brain-layer ΔHbO and Temple-BF traces (Fig. 5). Responses were averaged within predefined transition windows: −90 to 0s and 0 to 90s around transition onset for head-down tilt and stand-to-supine, and −45 to 0s and 0 to 45s for stand-to-squat. For each transition, the response was calculated as the mean value after transition onset minus the mean value before transition onset. Repeated baseline-to-task transitions within a protocol were then averaged within each session, giving one transition-response value per session, signal, and protocol.

**Fig. 5.**
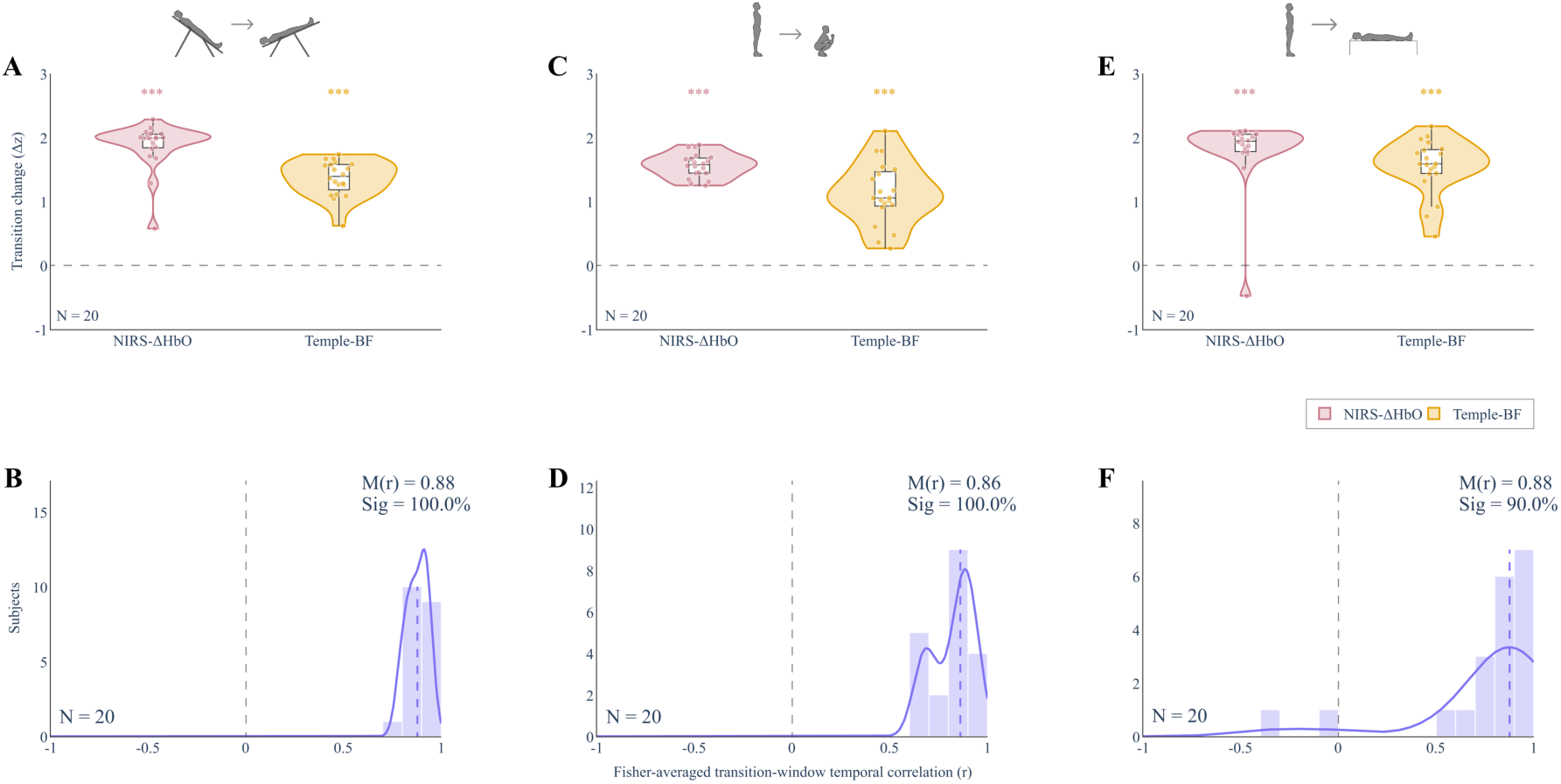
Directional concordance and transition correspondence between model-derived brain NIRS-ΔHbO and Temple-BF (n = 20 sessions per protocol). Baseline-to-task transition responses are shown for (A, B) head-down tilt, (C, D) stand-to-squat, and (E, F) the stand-to-supine protocol. Z-scored brain ΔHbO and Temple-BF traces were block-averaged within predefined transition windows. The top row shows the distribution of session-level transition changes for each signal; significance relative to zero was evaluated using a one-sided Wilcoxon signed-rank test, and “***” indicates p < 0.001. The bottom row shows the distribution of temporal correlations between the two z-scored signals within the corresponding transition windows. Correlations were calculated for each baseline-to-task transition window, and repeated transitions were combined within the session using Fisher r-to-z averaging. The median correlation, M(r), and Sig. are shown for each protocol. Sig is the percentage of sessions in which at least one repeated transition window showed a strong and statistically significant correlation, defined as r > 0.5 and p < 0.01.

Both brain-layer ΔHbO and Temple-BF increased during baseline-to-task transitions, with one-sided Wilcoxon signed-rank tests showing p < 0.001 for both signals across all protocols. Directional concordance was 100% for head-down tilt, 100% for stand-to-squat, and 95% for stand-to-supine, indicating that nearly all sessions showed transition-related changes in the same direction for the two signals.

After Fisher r-to-z averaging of repeated transitions within each session, median transition-window correlations were 0.878 for head-down tilt, 0.861 for stand-to-squat, and 0.876 for stand-to-supine. Group tests showed that these correlation distributions were significantly greater than zero (all p < 0.001). At the session level, at least one transition window met the criterion r > 0.5 and p < 0.01 in 100% of head-down tilt sessions, 100% of stand-to-squat sessions, and 90% of stand-to-supine sessions. Session-level results are summarized in Table 2.

**Table 2.**
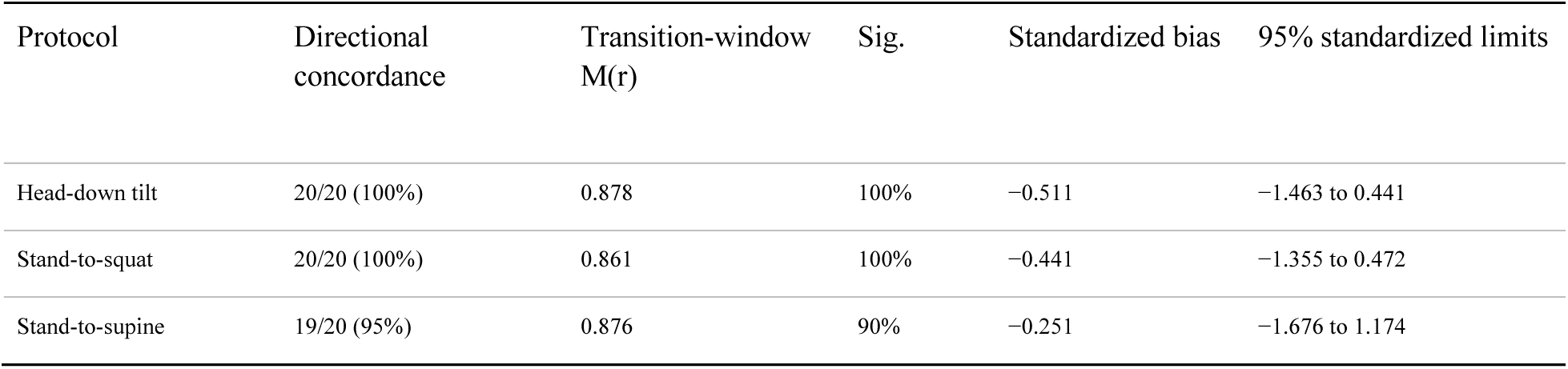
Transition-level comparison between Temple-BF and model-derived brain-layer NIRS ΔHbO by protocol (n = 20 sessions per protocol). Directional concordance is the number of sessions with transition responses of the same sign. Transition-window M(r) is the session-level median after Fisher r-to-z averaging of repeated baseline-to-task transition windows. Sig. is the percentage of sessions in which at least one repeated transition window met the criterion r > 0.5 and p < 0.01. Standardized bias and limits are calculated from z-scored transition-response magnitudes.

| Protocol | Directional concordance | Transition-window M(r) | Sig. | Standardized bias | 95% standardized limits |
| --- | --- | --- | --- | --- | --- |
| Head-down tilt | 20/20 (100%) | 0.878 | 100% | -0.511 | -1.463 to 0.441 |
| Stand-to-squat | 20/20 (100%) | 0.861 | 100% | -0.441 | -1.355 to 0.472 |
| Stand-to-supine | 19/20 (95%) | 0.876 | 90% | -0.251 | -1.676 to 1.174 |

Treating repeated transition windows separately (n = 40 windows per protocol; Fig. S5) gave similar median correlations of 0.888 for head-down tilt, 0.859 for stand-to-squat, and 0.846 for stand-to-supine. The percentage of individual transition windows meeting the criterion of r > 0.5 and p < 0.01 was 100.0%, 90.0%, and 85.0%, respectively.

### 3.6 Standardized paired-difference analysis

Bland-Altman plots were used to compare whether Temple-BF and brain-layer NIRS-ΔHbO captured similar transition-response magnitudes (Fig. 6). Because the two modalities have different native units, both signals were z-scored within session before calculating baseline-to-task transition changes. The analysis therefore evaluates agreement in standardized response magnitude, rather than agreement in absolute physical units. The plotted difference was defined as Temple-BF minus brain-layer NIRS-ΔHbO. Mean standardized differences were −0.511 for head-down tilt, −0.441 for stand-to-squat, and −0.251 for stand-to-supine, with 95% limits of agreement of −1.463 to 0.441, −1.355 to 0.472, and −1.676 to 1.174, respectively. The consistently negative bias indicates that the standardized Temple-BF transition response was smaller than the standardized brain-layer response, by roughly a quarter to a half of a within-session standard deviation, and the limits span approximately 1.8 to 2.9 standard deviations. Temple-BF therefore reproduced the direction and timing of the transition response more closely than its standardized magnitude.

**Fig. 6.**
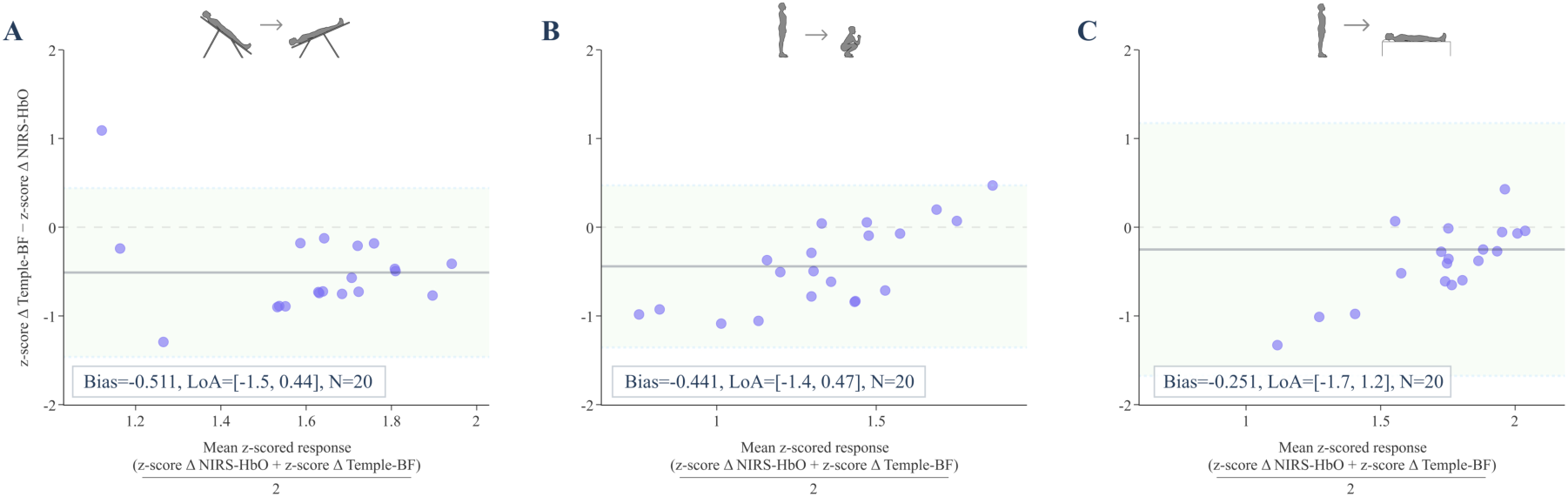
Bland-Altman plots of standardized transition responses from model-derived brain-layer NIRS-ΔHbO and Temple-BF (n = 20 sessions per protocol). Points show session-level paired transition responses after each signal was z-scored within-session. The x-axis is the mean of the two standardized transition responses and the y-axis is Temple-BF minus NIRS-ΔHbO. The solid line is the mean standardized difference, and dashed lines are mean ± 1.96 SD of the standardized differences.

## 4 Discussion

Across the three imposed paradigms, Temple-BF showed positive temporal association with the model-derived brain-layer ΔHbO signal, with median lag-adjusted correlations of 0.912, 0.843, and 0.839 and zero-lag medians of 0.826, 0.701, and 0.796 for head-down tilt, stand-to-squat, and stand-to-supine, respectively. Importantly, the zero-lag correlations remained substantial, supporting the interpretation that the Temple-BF and brain-layer ΔHbO association was not simply an artifact of the lag-search procedure. The direction of transition-related change matched in nearly all sessions, with 100% concordance for head-down tilt and stand-to-squat and 95% concordance for stand-to-supine. The broader physiological pattern is compatible with prior NIRS and cerebrovascular studies of tilt, squat-stand, and orthostatic transitions^28–30,48,49^. The principal finding is that Temple-BF preferentially tracked the deeper, brain-assigned compartment during large task-locked maneuvers, the pattern predicted if the index carries cerebrally relevant information.

### 4.1 The contrast between compartments

The brain-layer correlations exceeded the superficial-forehead in all three protocols, with the largest separation during head-down tilt protocol. This distinction is informative because head-down/head-up transitions alter hydrostatic pressure, venous pressure, cerebral perfusion pressure, and extracranial perfusion in ways that need not be identical across compartments^29,31,32^. Head-down tilt is a useful perturbation because intracranial and cutaneous vessels respond differently to the same hydrostatic perturbation. At −30° head-down tilt, intracranial pressure increases while mean middle cerebral artery velocity and dynamic autoregulation may remain relatively preserved^31^, whereas facial skin blood flow can decrease with increased cutaneous vascular resistance^32^. The latter observation is directionally consistent with our findings that scalp layer ΔHbT decreases. Though the inference is tentative because those measurements reflect blood flow rather than blood volume and were made at the cheek whereas our recordings were made over the forehead, and facial cutaneous vasomotor responses are not necessarily known to be uniform across regions^50^. Notably, the superficial response in our data was itself not uniform; ΔHbT decreased during head-down tilt despite an increase in superficial ΔHbO, implying a decrease in ΔHbR. Thus, the superficial measurements do not indicate a simple uniform reduction in cutaneous hemodynamics during head-down tilt. This contrasts with increases observed in Temple-BF and brain layer-derived ΔHbO and ΔHbT. But note that NIRS-based hemodynamic changes were estimated on the side of the forehead opposite the Temple sensor rather than directly beneath it.

### 4.2 Systemic and protocol-dependent differences

All three protocols evoke systemic as well as cerebral responses, including changes in arterial pressure, venous return, cardiac output, ventilation, CO₂, autonomic tone, and skin perfusion^27,33,44,51–54^. Adjusting for Temple-derived heart rate lowered the correlations for head-down tilt and stand-to-supine and slightly raised it for stand-to-squat, without eliminating the association in any protocol (Table 1). This shows only that linear heart rate variation does not account for the association, though it does not distinguish cerebral coupling from unmeasured systemic covariation. Arterial blood pressure is a plausible common input to both compartments: pressure fluctuations drive cerebral flow through frequency-dependent autoregulatory dynamics^55,56^, similar transfer-function methods have been applied to NIRS-derived cerebral hemodynamics^57,58^, and cutaneous vascular beds also exhibit distinct responses to systemic pressure perturbations^59^. The same pressure perturbation can therefore move both signals without producing identical time courses. Whether the Temple-BF algorithm preferentially retains pressure-coupled dynamics resembling the cerebral response cannot be determined without beat-to-beat pressure measurement.

Protocol-dependent effect sizes are physiologically plausible but not uniquely interpretable. The compartmental separation was most apparent during head-down tilt, where the median paired brain-minus-scalp difference was 0.33, compared with 0.11 for stand-to-squat and 0.04 for stand-to-supine (Table 1). Head-down tilt is passive and produces a large, slow stimulus with little voluntary movement, whereas squatting adds skeletal-muscle contraction, rapid hydrostatic change, possible movement artifact, and strong cardiovascular transients. Squat-stand maneuvers are widely used precisely because they impose large blood-pressure oscillations^27,33^. Stand-to-supine transitions also alter venous return and systemic cardiovascular state. These differences can change both true physiology and measurement artefacts. Because blood pressure, CO₂, respiration, and local superficial perfusion were not recorded, the protocol differences should not be attributed to a specific mechanism from the present dataset.

### 4.3 Assumptions in the two-layer inversion

The two-layer inversion depends on the structural and optical property assumptions stated in Sec. 2.4: a fixed 8mm superficial thickness, an assumed 1.5:1 brain-to-scalp baseline absorption ratio, and a two-layer slab geometry rather than individualized head anatomy. Time-domain moments can improve depth selectivity, but the recovered partition remains sensitive to geometry, optical properties, instrument response, and model conditioning^24–26,60^.

Screening poorly conditioned modules reduces numerical instability but does not make the compartment assignment assumption-free. This limitation was also evident when comparing the model-derived scalp estimate with the conventional short-SDS OD-derived HbO signal. The two-layer scalp trace did not always match the short-SDS OD-derived trace, even though both are intended to carry strong superficial sensitivity. One likely reason is that the scalp and brain sensitivity columns remain partially correlated in the two-layer inversion, so geometry or optical-property mismatch can redistribute shared signal between the modeled scalp and brain compartments. In this situation, some superficial temporal structure could be assigned to the brain layer, or conversely attenuated in the scalp layer, depending on the conditioning of the sensitivity matrix. This motivated the short-SDS partial-correlation sensitivity analysis, in which OD-derived HbO from channels with source-detector separations ≤9 mm was included as a superficial regressor. The persistence of positive brain-layer ΔHbO–Temple-BF partial correlations after this adjustment suggests that the Temple-BF association cannot be fully explained by superficial short-SDS HbO structure and shared association with the brain layer. But also, the deeper “brain layer” may include sensitivity to the meninges and pial vessels, rather than exclusively cortical microvasculature.

Temperature regression of mean TOF is a further assumption. It corrects thermal drift of the instrument response, but scalp and module temperature can change with posture^32^, so any task-locked component of the temperature trace is removed from TOF along with the drift. Because TOF carries the depth information in this inversion, this correction could attenuate genuine task-locked depth contrast as well as instrumental drift.

Variance of time of flight was available as a third observable but was excluded from the final inversion because its inclusion produced implausibly inflated hemoglobin amplitudes in several modules in this implementation. Variance in principle offers the highest depth selectivity among the moments^26^. But, the measured variance baseline was probably influenced by instrument-response broadening and possibly by light leakage at the optode interface, whereas the Monte Carlo sensitivities used here represented tissue-only variance changes. This mismatch between measured variance and tissue-only variance sensitivity caused the inversion to attribute non-tissue timing broadening to absorption changes, producing inflated hemoglobin amplitudes. A forward model that includes instrument response would be needed before variance can be used in this pipeline.

The 690/905nm wavelength pair also constrains chromophore separation. Tissue absorption at these wavelengths includes HbO, HbR, water and other chromophores^61^; the inversion modeled hemoglobin only, and this pair conditions ΔHbR more poorly than ΔHbO^42,62^. The weaker, protocol-dependent ΔHbR associations therefore should not be read as a physiological null. Since ΔHbO changes are relatively larger, closely similar associations with ΔHbO and ΔHbT are expected. Neither the Temple infrared nor the TD-NIRS pair is selective for a single chromophore^25,35,36^.

### 4.4 Limitations

1. The “brain” signal is a model-derived deeper compartment rather than a direct cortical measurement. Fixed scalp thickness, assumed baseline absorption contrast, a slab geometry, exclusion of TOF variance, and dataset-relative module screening can all influence compartment separation. Individualized anatomy and parameter-sensitivity analyses are needed to establish robustness.
2. The sample size was not determined by power analysis, maximum correlations were selected over a ±30 s lag search, one-sided tests and multiple secondary analyses were used without multiplicity adjustment. In addition, the common block design can produce high correlations when two sensors respond to the same imposed maneuver even if their tissue sources differ.
3. Systemic physiology was incompletely measured. Heart rate adjustment does not substitute for concurrent arterial pressure, end-tidal CO₂, respiration, SpO₂, cardiac output, or local superficial perfusion, all of which can influence NIRS and PPG during posture changes.
4. The superficial comparator was not colocated with the Temple sensor. TD-NIRS was recorded over the contralateral forehead while Temple-BF was recorded at the anterior temple. Because superficial hemodynamics are spatially heterogeneous, the forehead superficial estimate cannot exclude a local temple-scalp contribution.
5. Temple-BF is a proprietary relative index and surface PPG is fundamentally sensitive to local blood-volume changes in skin and subdermal vasculature^34,35^.
6. Long-term wear, test-retest reliability of Temple-BF within this protocol, contact-pressure effects, and sensitivity to smaller spontaneous changes typical of daily life were not established.
7. Participants were healthy men aged 20–32 years. Although Monk Skin Tone categories 3–8 were represented, the distribution by category and any association between pigmentation and signal quality or effect size were not reported. PPG performance can vary with biological characteristics, wavelength, sensor placement, temperature, and contact conditions, and single-sex recruitment limits generalizability to women, older adults, and clinical populations.

## 5 Conclusion

Across head-down tilt, stand-to-squat, and stand-to-supine challenges in healthy young men, Temple-BF showed positive temporal association with model-derived brain-layer TD-NIRS ΔHbO. Across all three maneuvers Temple-BF showed numerically larger median correlations with the deeper, brain compartment than with the concurrently estimated superficial scalp compartment. The separation was largest during head-down tilt, the condition that most strongly dissociates the two, and zero-lag analyses showed a similar pattern without lag optimization. These findings position Temple-BF as a potential relative marker of cerebral hemodynamic change during the postural perturbations evaluated in the study. Absolute flow accuracy, cerebral specificity at the temple, and the residual role of systemic and local superficial contributions remain to be established.

Future studies should combine the wearable with a flow-sensitive cerebral reference and comprehensive systemic physiology. Candidate designs include calibrated or quantitative perfusion imaging where feasible, diffuse correlation spectroscopy or transcranial Doppler for complementary flow/velocity information, and concurrent beat-to-beat arterial pressure, end-tidal CO₂, respiration, and SpO₂. A colocated short-separation or superficial optical channel at the temple would directly address local extracerebral contamination. Prespecified repeated-session testing should quantify within-subject reliability and smallest detectable change; individualized superficial thickness and sensitivity analyses should test the stability of the two-layer inversion. Recruitment should include women, broader age ranges, and prespecified analyses across skin pigmentation and hair/optode-coupling characteristics.

## Acknowledgments

The authors thank the study participants for their time and efforts, and Avi Roy for critical review of the manuscript.

## Code and Data Availability

De-identified TD-NIRS and Temple-BF time series supporting the findings of this article, together with the analysis code, are available from the corresponding author upon reasonable request. The algorithm generating the Temple Brain Flow Index is proprietary to Temple Private Limited and cannot be publicly released.

## Ethics approval

The study was approved by the Saarthak Ethical Research Institutional Ethics Committee (SER-IEC), New Delhi, India (approval SER-IEC/2026/AP/051). Written informed consent was obtained from all participants before any study procedure. This was an observational device-comparison study in healthy volunteers and not a clinical trial; no medical intervention, treatment, or randomization was involved.

## Appendix A: Supplementary figures

**Fig. S1.**
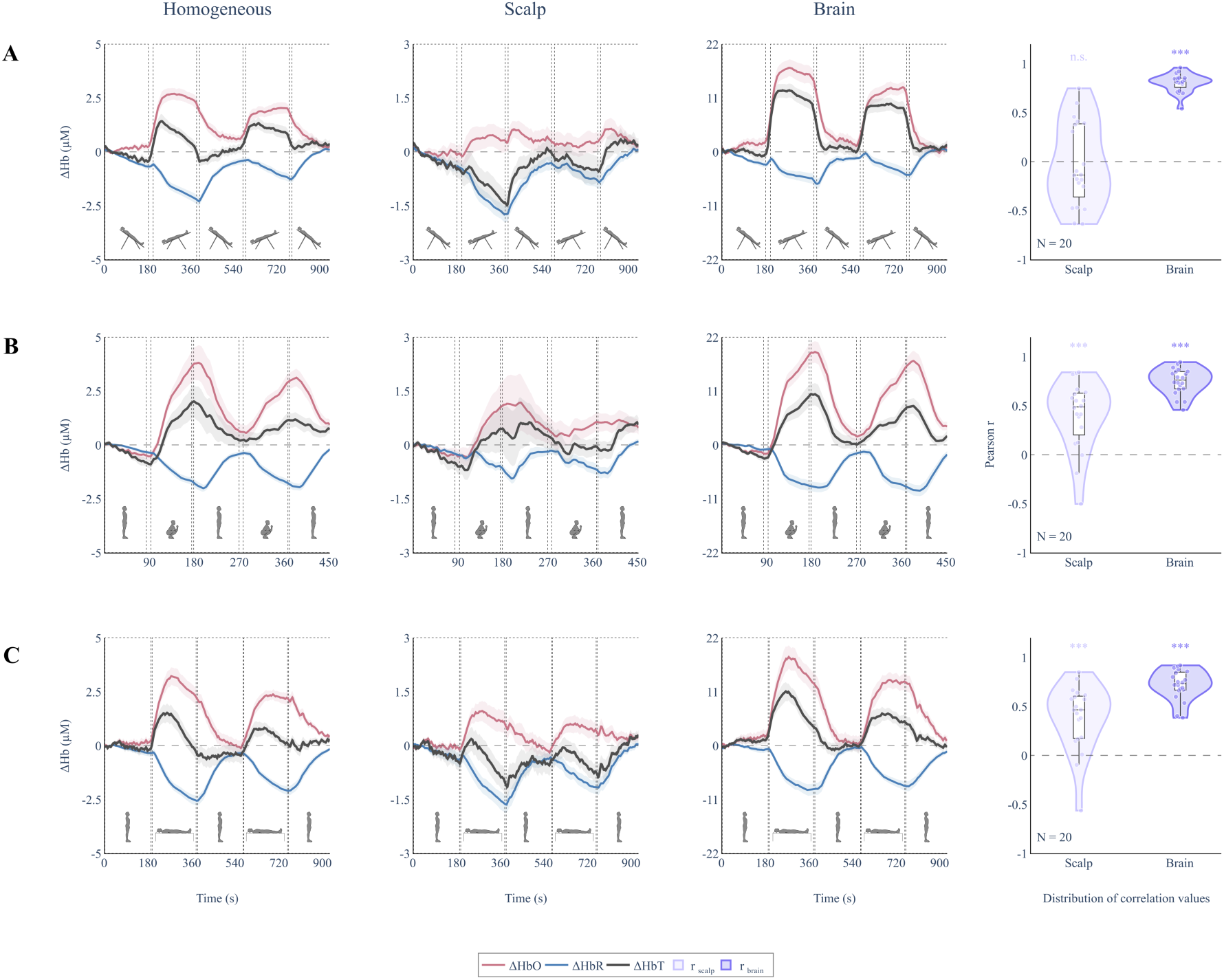
Model-derived NIRS hemoglobin responses with TDDR preprocessing. Model-derived NIRS hemoglobin responses across the three physiological challenge protocols using the same analysis pipeline as Fig. 3, but with TDDR applied to both OD and TOF before inversion. Group-averaged hemoglobin traces are shown for (A) head-down tilt, (B) stand-to-squat, and (C) the stand-to-supine protocol. Each row corresponds to one protocol, and the first three columns show the homogeneous estimate, scalp estimate, and brain estimate from the two-layer model. Traces show model-derived ΔHbO (pink), ΔHbR (blue), and ΔHbT (dark gray), with shaded bands representing SEM across sessions. Modules were screened using the same inversion-conditioning criteria as Figure 3, and only passing modules were averaged within each session. The fourth column shows the distribution of lagged Pearson correlations between Temple-BF and model-derived scalp or brain ΔHbO. Violin plots show session-level correlations, with embedded box plots and individual points.

**Fig. S2.**
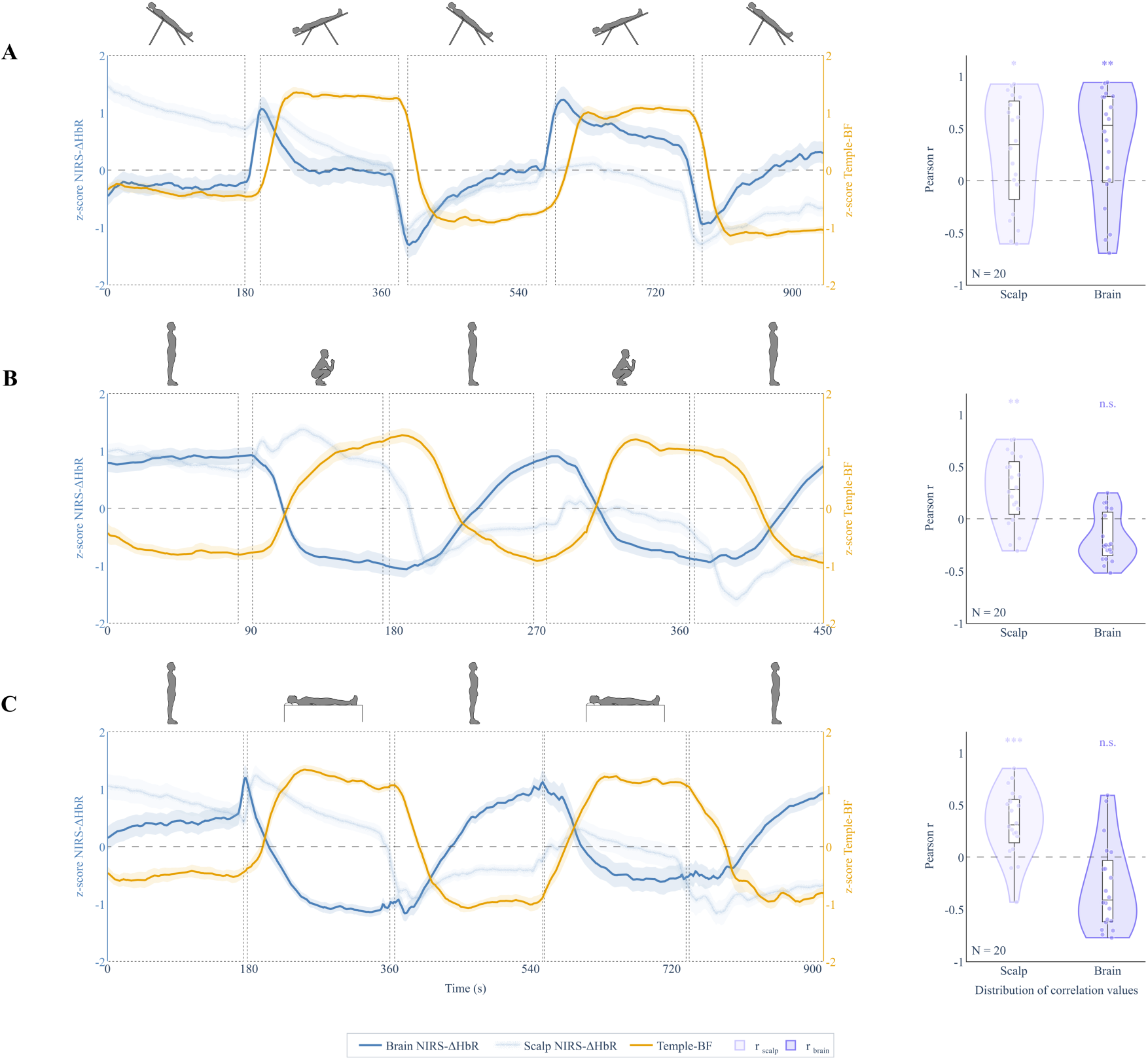
Temporal correspondence between model-derived NIRS-ΔHbR and Temple-BF across sessions and protocols. Same format as Fig. 4 but showing ΔHbR instead of ΔHbO. Brain-layer ΔHbR is shown as a solid blue trace, superficial ΔHbR as a light-blue trace, and Temple-BF in gold. Violin plots show session-level maximum lag-adjusted Pearson correlations.

**Fig. S3.**
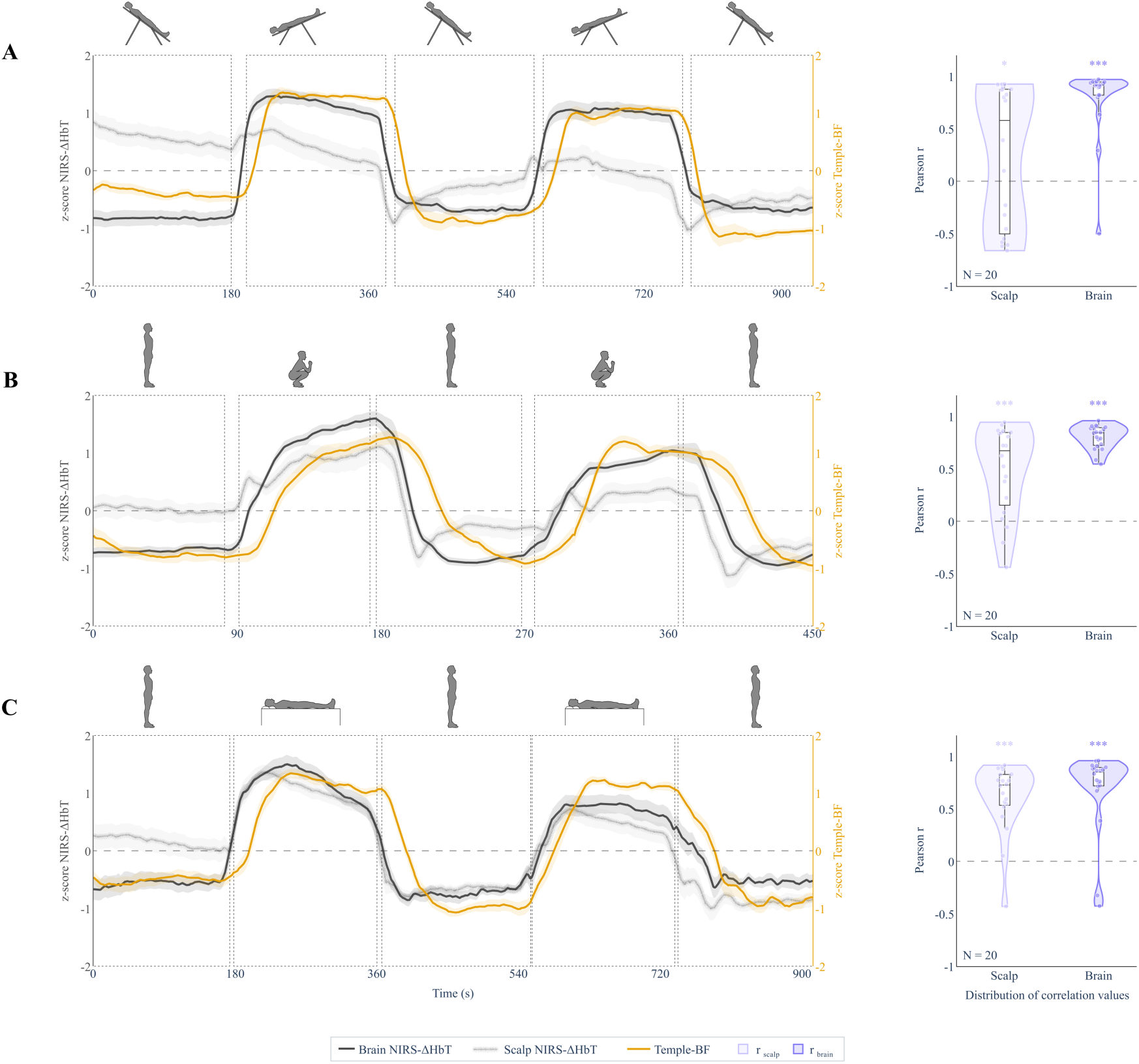
Temporal correspondence between model-derived NIRS-ΔHbT and Temple-BF across sessions and protocols. Same format as Fig. 4, but showing ΔHbT instead of ΔHbO. Brain-layer ΔHbT is shown as a solid dark-gray trace, superficial ΔHbT as a light-gray trace, and Temple-BF in gold. Violin plots show session-level maximum lag-adjusted Pearson correlations.

**Fig. S4.**
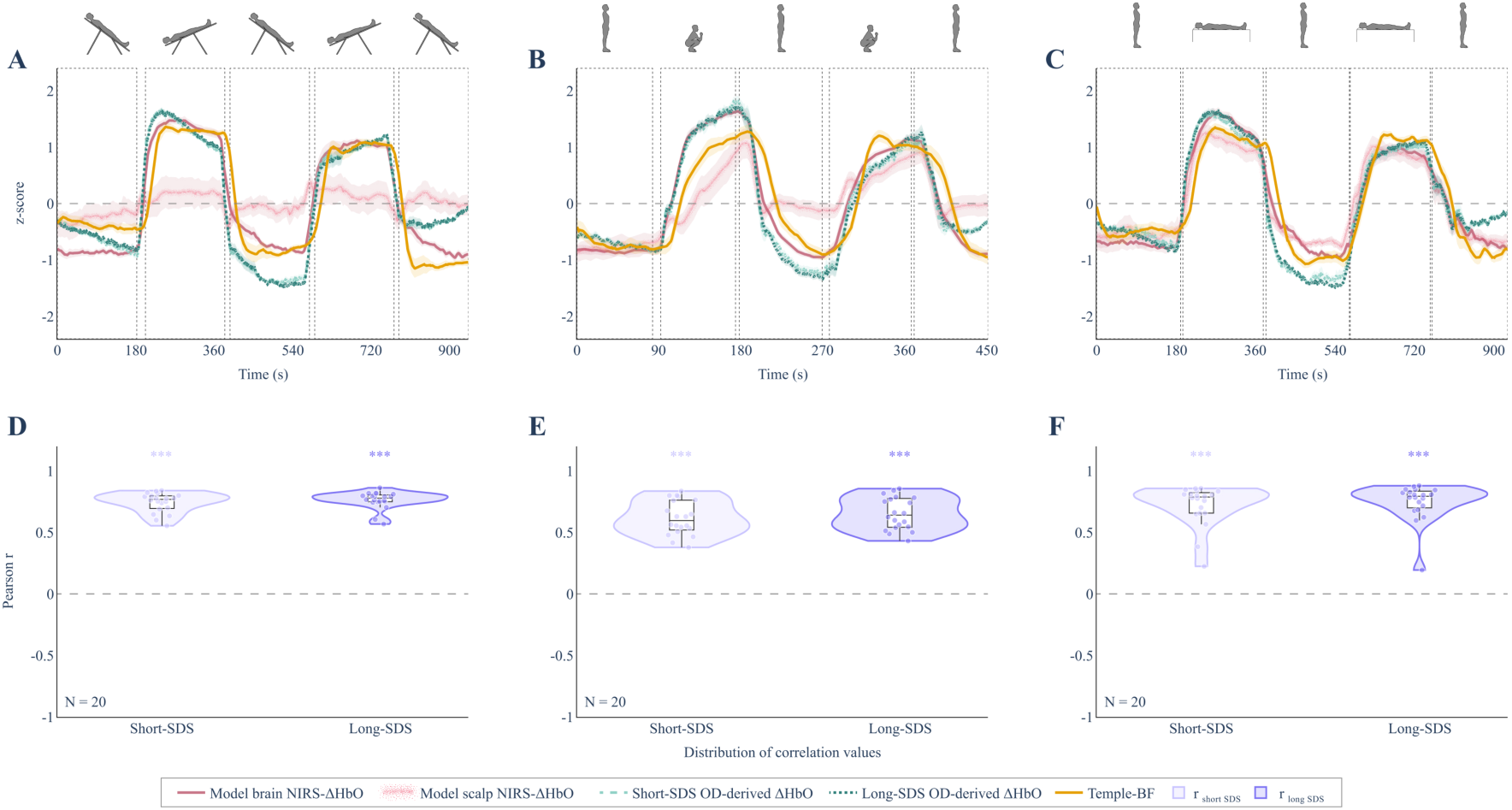
Model-derived and OD-derived HbO comparison with Temple-BF. Group-averaged z-scored time traces of model-derived brain NIRS-ΔHbO, model-derived scalp NIRS-ΔHbO, OD-derived short-SDS HbO, OD-derived long-SDS HbO, and Temple-BF are shown for (A) HDT, (B) stand-to-squat, (C) stand to supine protocol. Short-SDS and long-SDS OD-derived HbO were computed from channels with source-detector separations ≤9 mm and >9 mm, respectively. Dashed boxes indicate the experimental blocks. The lower row (D-F) shows the distribution of lagged Pearson correlation coefficients between Temple-BF and OD-derived HbO for short-SDS and long-SDS channels across participants. Violin plots include individual session values and embedded box plots. Asterisks indicate whether the correlation distribution was significantly greater than zero using a one-sided Wilcoxon signed-rank test.

**Fig. S5.**
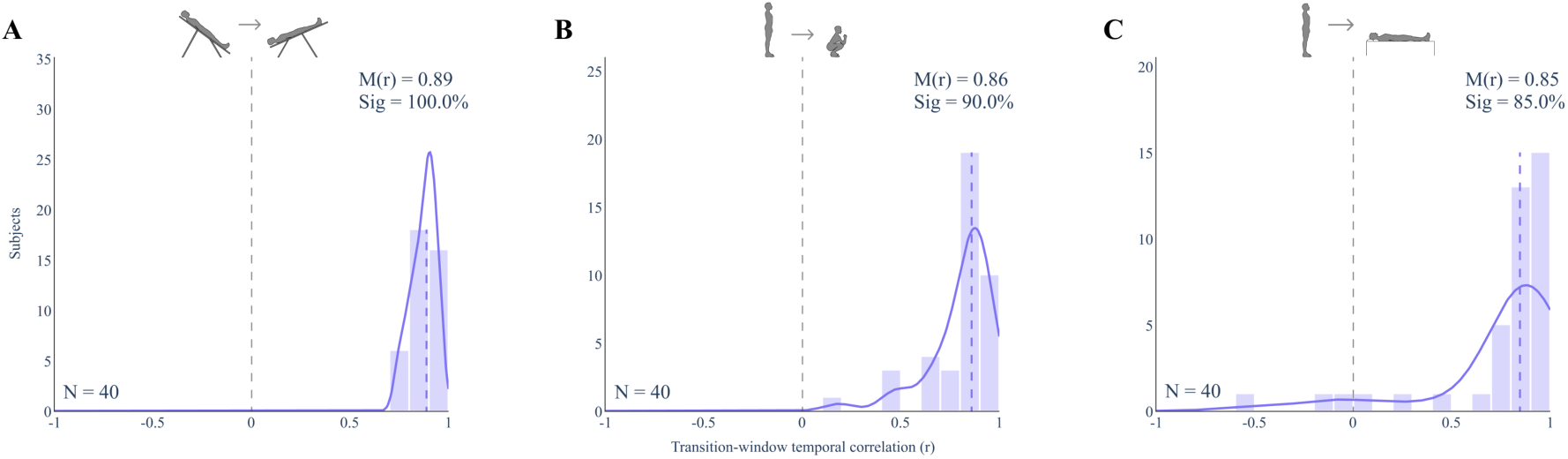
Transition-window temporal correspondence between model-derived brain-layer NIRS-ΔHbO and Temple-BF. Correlations are shown for individual baseline-to-task transition windows. Unlike Fig. 5, where repeated windows were combined within a session by Fisher r-to-z averaging, this figure shows all repeated windows separately (40 windows per protocol). M(r) is the median coefficient for each protocol. Sig. is the percentage of individual transition windows meeting r > 0.5 and p < 0.01.

